# Mode of sucrose delivery alters reward-related phasic dopamine signals in nucleus accumbens

**DOI:** 10.1101/132126

**Authors:** James E McCutcheon, Mitchell F Roitman

## Abstract

In studies of appetitive Pavlovian conditioning, rewards are often delivered to subjects in a manner that confounds several processes. For example, delivery of a sugar pellet to a rodent requires movement to collect the pellet and is associated with sensory stimuli such as the sight and sound of the pellet arrival. Thus, any neurochemical events occurring in proximity to the reward may be related to multiple coincident phenomena. We used fast-scan cyclic voltammetry in rats to compare nucleus accumbens dopamine responses to two different modes of delivery: sucrose pellets, which require goal-directed action for their collection and are associated with sensory stimuli, and intraoral infusions of sucrose, which are passively received and not associated with external stimuli. We found that when rewards were unpredicted both pellets and infusions evoked similar dopamine release. However, when rewards were predicted by distinct cues, greater dopamine release was evoked by pellet cues than infusion cues. Thus, dopamine responses to pellets, infusions as well as predictive cues suggest a nuanced role for dopamine in both reward seeking and reward evaluation.

## Introduction

The mode in which a reward is delivered may affect how it is perceived and encoded by neural circuits. First, different modes of reward delivery may affect the subsequent behaviour required to receive reward. For example, if food rewards – either in pellet or liquid form – are delivered to a receptacle animals need to attend to the delivery and organize approach behaviour before the food can be consumed. In contrast, solutions delivered through an intraoral cannula directly into the oral cavity are immediately available for consumption without requiring effort or movement (Grill and Norgren, 1978). Second, the sensory processes that are engaged by different modes of delivery will differ between foods in a solid or liquid form. Finally, the rate at which a reward is ingested or absorbed may affect its rewarding or reinforcing properties and the neural processes that are engaged (Avena et al., 2008; Ferrario et al., 2008; Furlong et al., 2014; Samaha et al., 2002). All of these factors may influence both how reward delivery and receipt are encoded by the brain and how information about associated stimuli and contexts is processed.

Dopamine signaling in nucleus accumbens (NAc) has been strongly associated with responses to reward and reward-related stimuli (Bassareo and Di Chiara, 1997; Gunaydin et al., 2014; Roitman et al., 2004; Steinberg et al., 2013; Stuber et al., 2005). Particularly important are brief, phasic, increases in concentration as seen in NAc following unpredicted presentation of rewards, such as food pellets, as well as after presentation of initially neutral stimuli (cues) that become reliable predictors of reward (Brown et al., 2011; Day et al., 2007; Flagel et al., 2011). These studies - of dopamine release in terminal regions - have predominantly been conducted in rodents using sugar pellets or sugar solution delivered to a receptacle, and therefore require the organization of appetitive behaviors for collection. Another less commonly used mode of reward delivery is intraoral infusions, which via an implanted cannula, allow solutions to be delivered directly to the oral cavity (Grill and Norgren, 1978). This technique is favored in certain situations as it allows exquisite experimenter control over stimulus exposure and may allow the consummatory phase of ingestive behaviour to be isolated from appetitive behaviors (Hudson and Ritter, 2004; Seeley et al., 1995). A small number of studies have examined phasic dopamine responses to unpredicted intraoral sucrose infusions with most evidence showing, relative to pellets, responses to the infusions themselves are smaller and less phasic in nature (McCutcheon et al., 2012; Roitman et al., 2008). However, to our knowledge, dopamine responses to pellets and infusions have never been directly compared and, furthermore, responses to cues that predict intraoral sucrose infusions have not been examined (see Cone et al., 2016 for responses to sodium infusion-paired cues).

Here, we have directly compared phasic dopamine responses in NAc to sucrose pellets and intraoral infusions in the same rats and, importantly, within the same dopamine recording session. In addition, all rats were trained to associate distinct predictive cues with receipt of each mode of sucrose delivery and dopamine responses to each cue were compared. Finally, due to the proposed functional heterogeneity of striatal subregions (Kelley, 1999) and identification in previous reports of regional differences in responses to rewards and predictive cues (Brown et al., 2011; Cacciapaglia et al., 2012; Wheeler et al., 2011), we compared responses across NAc core and shell.

## Results and Discussion

### Training and testing schedule

Rats were trained in a Pavlovian conditioning paradigm in which distinct cues (spatially distinct lever-cue light combinations) reliably predicted delivery of either a sugar pellet into a food receptacle or an infusion of the same amount of intraoral sucrose. After seven conditioning sessions in which rats experienced approximately thirty of each cue-reward pairing per session, all rats underwent a recording session in which cued and uncued rewards were presented in a pseudorandom order while dopamine was measured in NAc (see Methods for detailed information).

### Phasic dopamine signals during uncued trials

To assess whether different patterns of dopamine release were associated with the different modes of sucrose delivery (trial types), we used fast-scan cyclic voltammetry to measure changes in dopamine concentration during behaviour (Fig. 1). During the test session, uncued trials were interleaved with trials in which each reward was preceded by a distinct, predictive cue. Representative examples of dopamine responses to each of these trial types are shown in Fig. 1A and B.

**Figure 1.**
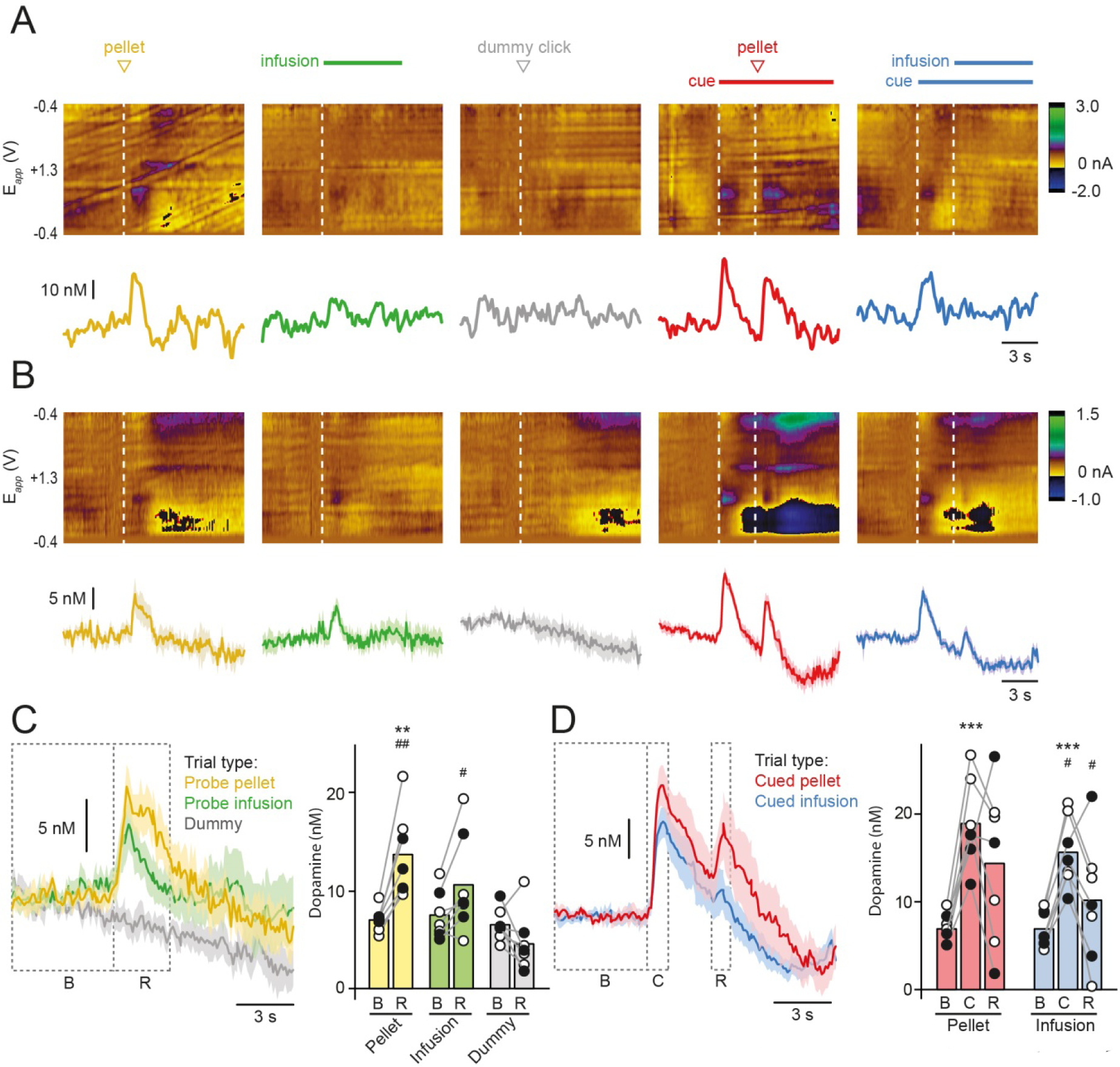
Dopamine responses to cues and primary rewards are qualitatively similar despite different types of reward delivery. **A,** Representative fast-scan cyclic voltammetry data from a single rat (electrode in NAc core) showing single trial examples of each trial type. Upper panels show colour plots with time shown on the x-axis, electrode holding potential shown on the y-axis, and background-subtracted current shown in pseudocolour. Vertical dashed white lines show events. Dopamine concentration, extracted using principal component analysis, is shown in lower panels. **B,** Averaged colour plots derived for each trial type for the same rat shown in A. Mean dopamine concentration traces ± SEM are shown below. **C,** Dopamine response averaged from all rats (core and shell) for cued trials (left) and background-subtracted dopamine concentration during baseline (B), cue presentation (C), and reward delivery (R) epochs (right). Bars show mean and circles show data from individual rats with core rats represented by empty circles and shell rats as filled circles. **D,** Average dopamine concentration data from all rats for probe trials in which no cue was presented (left) and background-subtracted dopamine concentration during baseline (B) and reward delivery (R) epochs. Bars show mean and circles show data from individual rats as coded in C. ***, p<0.001; **, p<0.05 vs. baseline epoch. ##, p<0.01, #, p<0.05 vs. corresponding epoch during dummy trials in C and pellet trials in D.

First, we compared trials in which pellets and infusions were delivered without a predictive cue to examine whether, in this situation, each type of reward was associated with a differential dopamine response (Fig. 1C). In addition, we included trials in which a dummy solenoid click was used but no infusion was delivered to ensure that any responses to the infusion were not a conditioned response to the potentially audible opening of the valve (importantly, both the real solenoid and dummy solenoid were housed outside a large sound-insulated chamber and should be inaudible to the rat). To assess dopamine responses across the extent of NAc, in these initial analyses we pooled data from both core and shell subregions. Two-way within-subjects repeated measures ANOVA revealed significant main effects of Trial Type (F(2,12)=13.689, p=0.001) and Epoch (F(1,12)=8.895, p=0.025), as well as a significant Epoch x Trial Type interaction (F(2,12)=23.139, p<0.001). Further testing revealed that on pellet trials dopamine was significantly elevated above baseline at time of reward delivery (p=0.006), and there was a trend for elevated dopamine during reward delivery on infusion trials (p=0.082). On dummy trials no dopamine responses were seen, relative to baseline (p=0.404). In addition, analyzing each epoch separately revealed that dopamine was similar during the baseline epoch for all trial types (F(2,12)=0.932, p=0.421) whereas, during the reward epoch, there was modulation by trial type (F(2,12)=20.132, p<0.000). *Post hoc* tests revealed that, relative to dummy trials, dopamine was elevated by both pellets (p=0.002) and infusions (p=0.011) whereas the concentration of dopamine evoked by pellets and infusions did not differ (p=0.310). Thus, when rewards were uncued, delivery of both pellets and infusions evoked an increase in dopamine concentration of similar magnitude.

### Phasic dopamine signals during cued trials

Next, we examined trials in which Pavlovian cues were included to assess whether there was differential dopamine release evoked by the cues based on the predicted mode of sucrose delivery. In addition, we examined how the presence of a predictive cue altered dopamine signaling at the time of reward delivery. To determine whether rats had learned to discriminate between the cues, we monitored the proportion of cued trials on which rats made a nose poke into the pellet receptacle. On test day, during the pellet cue rats made more nose pokes into the receptacle than during the infusion cue (Fig. 2A; p=0.028, n=7). In addition, when we considered time spent nose-poking in the pellet receptacle during presentation of each cue, but before reward delivery, we found that the pellet-predictive cue elicited a greater amount of time spent nose-poking in the pellet receptacle, relative to the infusion-predictive cue (pellet cue, 1.93 ± 0.26 s; infusion cue, 1.54 ± 0.23 s; t(6)=3.767, p=0.009).

**Figure 2.**
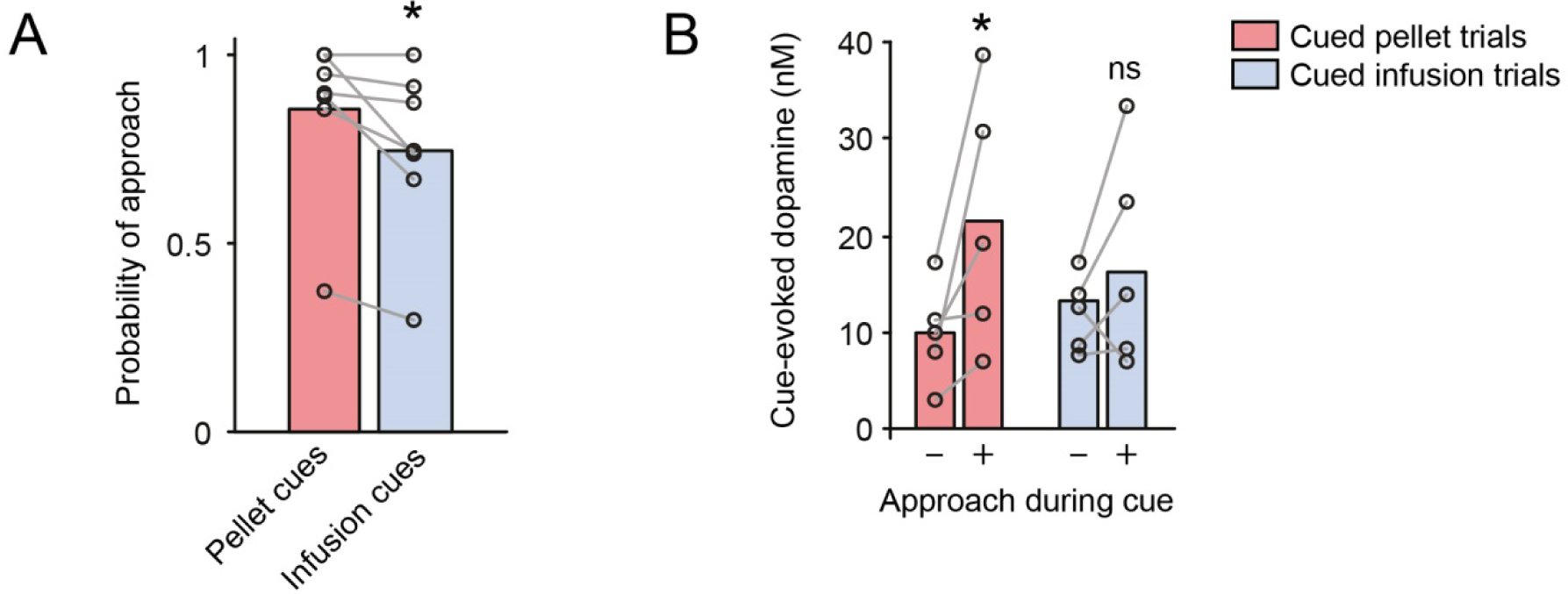
Cue-evoked dopamine responses discriminate between trials with approach and trials without approach only for pellet-predictive cues. **A,** Probability of performing a nose-poke in the pellet receptacle during cue presentation is greater for pellet-predictive cues than infusion-predictive cues. **B,** When cues predict pellets, cue-evoked dopamine is greater during trials on which rats perform pose-pokes during the cue vs. trials on which they do not. In contrast, this difference in dopamine levels is not seen for cues that predict infusion. *, p<0.05 vs. pellet cues in A and vs. trials with no approach during cue in B.

Comparison of phasic dopamine signals pooled from NAc core and shell during these cued trials showed that mode of reward delivery influenced the dopamine response to cues and rewards (Fig. 1D). Two-way within-subjects repeated measures ANOVA revealed significant main effects of both Trial Type (F(1,12)=17.841; p=0.006) and Epoch (F(2,12)=12.202; p=0.001) with a significant Trial Type x Epoch interaction (F(2,12)=6.754; p=0.011). For cued pellet trials there was a significant modulation of the dopamine concentration across epochs (F(2,12)=11.566, p=0.002) with *post hoc* tests revealing that, relative to baseline, there was elevated dopamine concentration during the cue epoch (p<0.001) and a trend for an elevation during the reward epoch (p=0.084). For cued infusion trials, a significant modulation of dopamine was seen across epochs (F(2,12)=12.206, p=0.001). However, although dopamine was elevated, relative to baseline, during the cue epoch (p<0.001) dopamine was not elevated during infusion delivery (p=0.368). Direct comparison of dopamine concentration during each epoch in pellet and infusion trials revealed that there was no difference in concentration during baseline epoch (p=0.999) but that, in pellet trials relative to infusion trials, dopamine was elevated in both the cue epoch (p=0.033) and the reward epoch (p=0.042).

To assess whether this difference in dopamine during the cue epoch was associated with a difference in behaviour, we compared cue-evoked dopamine on trials in which rats made a nose poke during the cue vs. trials on which rats did not. This analysis revealed that during the pellet-predictive cue, cue-evoked dopamine was elevated on trials where rats approached vs. trials on which they did not (Fig. 2B, red bars; p=0.043, n=5). In contrast, on infusion trials, cue-evoked dopamine was not different on trials with approach vs. trials without approach (Fig. 2B, blue bars; p=0.463, n=6). Taken together, these analyses show that both cues evoked similar patterns of dopamine release, however, the magnitude of this release was greater for rewards that require an additional response (e.g. approach to consume sucrose pellets) vs. passively-received rewards (sucrose infusions). In addition, while delivery of pellet rewards increased dopamine, delivery of infusions did not.

### Subtle differences are seen between phasic dopamine release in NAc subregions

Finally, we examined whether the pattern of responses differed across NAc subregions as there is a substantial body of work indicating that core and shell may encode different aspects of reward-related tasks (Kelley, 1999). Histological examination of lesion sites confirmed that recordings from four rats were made in NAc core and three rats were made in NAc shell (Fig. 3A). We binned data from each trial type into 500 ms bins and used receiver-operator characteristic (ROC) analysis to ask if dopamine responses in core and shell differed significantly at any time point (Fig. 3B and C). We found that, for each trial type, subtle differences existed between core and shell subregions (red circles on Fig. 3C show time points at which ROC analysis produced *p* < 0.05).

**Figure 3.**
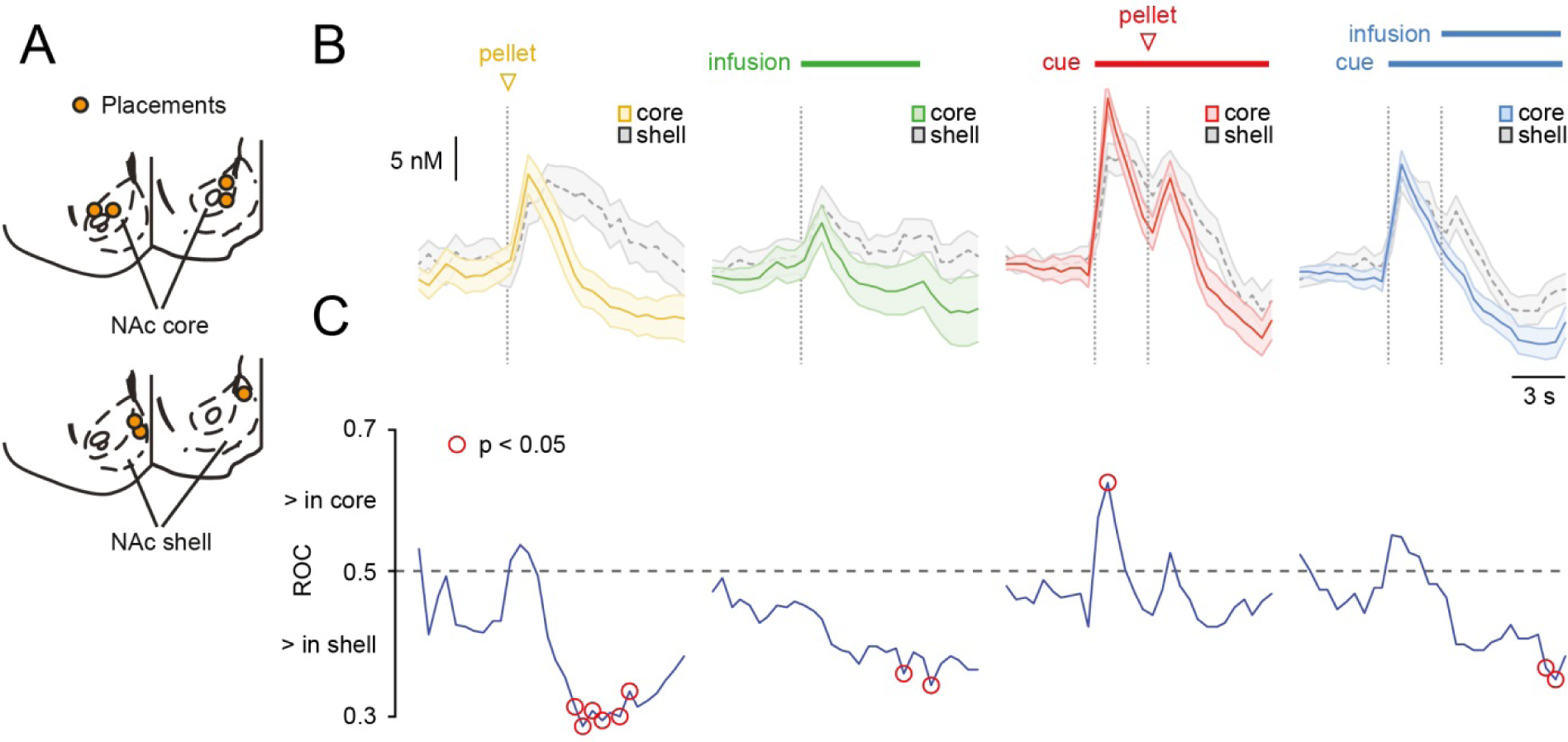
Dopamine responses in nucleus accumbens (NAc) core and shell show subtle differences in time course but not in principal features. **A,** Schematics to show electrode placements (yellow circles) in coronal sections at +2.0 mm (left) and +1.7 mm (right) from Bregma. **B,** Dopamine concentration traces evoked by different trial types in NAc core (solid lines, coloured shading) and shell (dashed lines, grey shading). **B,** Receiver-operator characteristic (ROC) analysis to determine time points at which core and shell responses deviate ROC values greater than 0.5 indicate elevated dopamine in core vs. shell and ROC values less than 0.5 indicate elevated dopamine in shell vs. core. Red circles denote time points at which this difference is statistically significant (p<0.05, Bonferroni corrected for multiple comparisons). Although the shapes of each response are similar in both regions, responses are more prolonged in shell following reward delivery, for example elevated dopamine seen in shell vs. core after reward delivery in uncued pellet trials, uncued infusion trials, and cued infusion trials. In addition, in cued pellet trials, cue-evoked peaks are sharper in core than in shell leading to elevated dopamine at time of cue.

In trials with uncued pellets or infusions as well as trials with cued infusions, a prolonged reward-evoked elevation of dopamine was seen in shell, relative to core, that was apparent several seconds after reward delivery. An elevation of dopamine in the core, relative to the shell, was only observed on trials with cued pellets and this elevation was seen immediately following cue presentation. Taken together, these results show, surprisingly, that in this paradigm only fairly subtle differences exist in the dopamine responses to cues and rewards across subregions.

### Conclusions and Discussion

We demonstrate that during presentation of uncued rewards, a requirement for goal-directed action had little effect on the dopamine response to reward as both pellets and infusions evoked a similar increase in dopamine concentration. When predictive cues were present, subtle differences emerged. Although cue-evoked dopamine was qualitatively similar in both pellet and infusion trials, dopamine evoked by pellet cues was of greater magnitude than that evoked by infusion cues. In addition, when rewards were cued, ‘primary’ reward evoked greater dopamine release on cued pellet trials than on cued infusion trials. These results suggest that dopamine’s role in signaling reward prediction and reward evaluation is nuanced and partially influenced by the mode of reward delivery.

Recent work suggests a tight link between high concentration, phasic increases in dopamine and the engagement in proactive behaviors for the consumption of food reward (du Hoffmann and Nicola, 2014; Hamid et al., 2015; Syed et al., 2015). Consistent with this framework, phasic dopamine signals emerge during Pavlovian conditioning along with conditioned approach (Aragona et al., 2009; Cone et al., 2016; Day et al., 2007; Stuber et al., 2008). The demonstration here that cues predicting passively-received intraoral infusions evoke dopamine is important because in most studies linking dopamine fluctuations to appetitive Pavlovian learning, approach behaviour of some kind has been required to retrieve the reward. The present study is the first to assess rewards matched for caloric content (i.e. sugar pellets vs. sugar solution) but that require distinct behavioral responses so that the magnitude and dynamics of dopamine release can be compared between cases. Thus, the finding that cues predicting intraoral delivery evoke dopamine is consistent with a role for dopamine in reward prediction independent of the action the cues instruct. Although work in monkeys recording somatic action potentials in dopamine neurons has often used behavioural paradigms in which rewards were presented within licking distance (Bromberg-Martin and Hikosaka, 2009; Schultz et al., 1993), this study is the first, to our knowledge, where dopamine release in terminal regions has been recorded during a task that requires no movement to retrieve rewards. We have recently shown that cues that predict intraoral salt differentially evoke NAc dopamine depending on physiological state (sodium appetite; Cone et al., 2016). The present study extends these findings by showing that intraoral sucrose rewards behave similarly and lead to cue-evoked dopamine responses. Finally, it remains possible that both cue types lead to enhanced incentive salience and thus equally recruit dopamine signaling (Berridge and Robinson, 2016).

Despite this demonstration that a goal-directed action was not required to receive reward on infusion trials, our data do not rule out a contribution of motor generation to dopamine signals. Indeed, goal-directed action could underlie the difference in magnitude that we observed between pellet-predictive and infusion-predictive cues. In our paradigm, although rats did not need to move to acquire intraoral infusions, they were also not compelled to stay still and, although the rats made fewer entries into the food receptacle on infusion trials than on pellet trials, they performed head entries on many trials nonetheless. This likely reflects either generalization of the cue-reward contingency, incomplete learning, and/or the fact that inappropriate head entries were not punished. Different training conditions may be needed to disentangle these possibilities such as a paradigm where errors are punished with timeouts or withholding of reward. For example, in a Go-NoGo paradigm when rats are required to suppress all movement and remain still during presentation of a reward-predictive cue, suppression of dopamine in NAc core is observed (Syed et al., 2015). In a different paradigm, under a Go-NoGo schedule in which rats must withhold prepotent responding on a lever to avoid a timeout, differential activity in NAc neurons is observed (Roitman and Loriaux, 2014).

Although we tried to match rewards in terms of their value to the rats, there were differences between rewards that may have affected our results. For example, pellet delivery involves additional audiovisual elements not present during intraoral delivery (e.g. magazine turn, rattling of the pellet down the chute and into the receptacle). For intraoral delivery, we explicitly masked any proximal cues that can be associated with these infusions such as the click of the solenoid valve. Thus, dopamine evoked at time of reward delivery in pellet trials may reflect cue-driven processes as well as goal-directed action needed to retrieve pellets. Another potential difference between trial types that could drive differential dopamine responses is the time course of reward receipt. Although rewards were matched for quantity/caloric value, in the case of infusions, delivery of sucrose spanned several seconds whereas, for pellets, rats received the sucrose at a single point in time and then, presumably, took a few seconds to chew and swallow the pellet. These differences between modes of reward delivery may influence the ability of rats to learn cue-reward associations that in turn affect how these stimuli and associations are encoded by dopamine and other brain structures.

With respect to dopamine signals at time of reward, it was of interest that in cued trials an dopamine was increased during delivery of pellets, relative to infusions. Interestingly, during uncued trials we saw no difference between the amount of dopamine evoked by pellets or infusions. Thus, a simple explanation involving reduced detection ability of infusions, relative to pellets, is ruled out. One potential explanation that lends support to a dual-encoding hypothesis is that, in the uncued situation, increases in dopamine subserve primarily a ‘reward-predictive’ role; both pellets and infusions are equally unexpected and so similar signals are seen. In cued trials, classic reward prediction error theory would suggest that no signal should be present at time of reward if the cue is completely predictive (Schultz, 1998). This is indeed what is observed on infusion trials. However, on pellet trials when additional action is required to retrieve reward, a robust increase in dopamine is observed, which may be necessary to invigorate retrieval (Nicola, 2016). An alternative explanation is that there are differences in the rate of learning for each cue-reward association. As such, it is possible that the cue-reward association develops more rapidly on infusion trials than on pellet trials, potentially because of the tighter temporal coincidence between events (e.g. infusions are always received exactly three seconds after cues whereas pellet receipt may be more delayed especially in early training sessions). Therefore, the predictive power of the infusion cue may be stronger than the pellet cue leading to a reduced dopamine response to cued infusions, relative to cued pellets. Longitudinal recordings from NAc across training sessions will help to disentangle these possibilities.

Phasic dopamine signals in NAc core and shell subregions may reflect different aspects of reward (Brown et al., 2011; Cacciapaglia et al., 2012; Saddoris et al., 2015; Wheeler et al., 2011). In particular, in naïve rats, dopamine in NAc shell responds to delivery of rewarding intraoral stimuli (McCutcheon et al., 2012; Roitman et al., 2008; Wheeler et al., 2011) whereas dopamine in NAc core appears unresponsive (Wheeler et al. 2011). In contrast, in trained rats, responses to food pellets or food-associated cues are seen in NAc core but not in NAc shell (Brown et al., 2011). Thus, it has been proposed that NAc shell is associated with responses to primary reward and that NAc core is more important for signaling environmental associations consistent with a later role in prediction error learning (Aragona et al., 2009). This framework follows an influential hypothesis involving the spiraling of information from ventromedial to dorsolateral parts of striatum during associative learning (Everitt and Robbins, 2005; Haber et al., 2000; Willuhn et al., 2012). Based on these previous findings we expected to observe clear differences in the current study between core and shell. Specifically, our hypothesis was that responses to cues would only be seen in NAc core and that NAc shell would only respond to ‘primary’ rewards. Surprisingly, this distinction was not present in our data; instead we found that NAc core and shell responses to each event were, in general, remarkably similar. We did observe differences in the time course of dopamine release events: responses tended to be more prolonged in the shell, relative to the core. The latter pattern could reflect the known differences in dopamine transporter expression between these regions; low levels of expression in the shell allow dopamine responses to be persist for longer than in the core (Ciliax et al., 1995). Reasons for the discrepancy between the findings here and other studies may reflect the different tasks animals were engaged in, the level of training animals received, or even different rewards under study. One hypothesis is that, in the present study, the (relatively) low level of training and competing cue-reward associations may have kept NAc shell online and engaged in processing stimuli. In fact, it is not unprecedented to find responses to cues in NAc shell in studies using self-paced operant conditioning and more complex tasks (Cacciapaglia et al., 2012; Saddoris et al., 2015).

To summarize, different modes of reward delivery allow the appetitive and consummatory phases of ingestive behaviour to be studied independently. Here, we used a combination of cued and uncued intraoral infusions and sucrose pellets to assess how dopamine encodes multiple events in the sequence of actions that ultimately produce feeding. Our results demonstrate that, in line with recent theoretical work (Berke, 2018), dopamine has a broad, nuanced role that does not seem restricted to either phase of behaviour but rather may contribute to goal-directed action and motivation, reward prediction, and stimulus evaluation.

## Methods

### Subjects

Male Sprague-Dawley rats (n=7; Charles River) weighing 325-375 g and aged approximately 10-12 weeks at the start of the experiment were used. Rats were individually housed with lights on from 7:00 to 19:00. All rats were food-restricted to 90-95% of free-feeding weight for 2-3 days before behavioural training began. Food restriction continued throughout the experiment with the exception of approximately 2 days before and 7 days after surgical procedures when rats were provided with ad libitum food. Animal care and use was in accordance with the National Institutes for Health Guide for the Care and Use of Laboratory Animals, and approved by the Institutional Animal Care and Use Committee at the University of Illinois at Chicago.

### Surgical procedures

Procedures were as described elsewhere (Cone et al., 2016; Fortin et al., 2015). Briefly, under general anesthesia (100 mg/kg ketamine + 10 mg/kg xylazine; i.p.), rats were implanted with an intraoral cannula in a first surgery and, after initial training (see below), were implanted with apparatus for performing fast-scan cyclic voltammetry recordings in a second surgery. This apparatus consisted of: a guide cannula (Bioanalytical Systems; West Lafayette, IN) directed towards NAc core (mm from Bregma: +1.3 AP, +1.3 ML; n=4) or shell (mm from Bregma: +1.7, +0.9 ML; n=3) and a Ag/AgCl reference electrode in contralateral cortex secured to the skull using stainless steel screws and dental cement. Rats were given meloxicam (1 mg/kg) and enrofloxacin (10 mg/kg) at time of surgery and for two days post-operatively. Rats had at least 1 week of recovery between each surgery and continuation of the experiment.

### Behavioural procedures

Behavioural training, testing, and fast-scan cyclic voltammetry experiments took place in the same chambers (20 cm length x 12 cm width x 14 cm height) equipped with: a pellet receptacle; a house light; a white noise generator; two levers and two cue lights flanking the pellet receptacle; and a fluid line attached to a solenoid valve and solution reservoir. The solenoid valve was positioned outside of the behavioural chamber to minimize the likelihood that rats could use the audible click of the valve as a predictor of sucrose delivery. In all sessions the house light was illuminated and the white noise generator was applied (60 dB). Two different reward stimuli, matched for caloric content, were used: 45 mg sucrose pellets (BioServ) and ~329 µL intraoral infusions of 0.4 M sucrose solution. Intraoral infusions (6.5 s duration) were delivered by activating the solenoid valve to allow sucrose solution to flow directly into the rats’ mouth at a rate of ~50 µL/s. Rats were given 2-6 pre-testing sessions to become familiar with each type of reward. First, rats were presented with a session in which 30 sucrose pellets were delivered to the receptacle at pseudorandom intervals (mean ITI 45 s; range 30 – 60 s). The number of pellets consumed was recorded and the following day, each rat was placed back into the chamber and received the same number of infusions in a similar temporal pattern to match pellet consumption from the previous day. This pattern (pellet day, infusion day) was continued until rats consumed all pellets delivered, typically 2-3 repetitions. Once rats reached criteria, they received one day of training in which pellets and infusions were delivered at pseudorandom intervals and in pseudorandom order. Following this session, Pavlovian conditioning sessions began. In these sessions, delivery of each reward was preceded by extension of a lever and illumination of the cue light above the lever (e.g. right lever and cue light → sucrose pellet vs. left lever and light → sucrose infusion; sides counterbalanced across rats). Importantly, interaction with the levers had no programmed consequence. Each reward was delivered 3 s after the cue onset and the cue remained on until 6.5 s after reward delivery, the time at which the infusion stopped. An infrared beam was situated above the pellet receptacle allowing head entries into the receptacle to be recorded. For training sessions, rats were presented with 28-30 trials of each cue-reward pair in a pseudorandom manner. Rats experienced five of these sessions before undergoing surgery for fast-scan cyclic voltammetry. Following recovery from surgery, rats experienced 1-2 post-surgical training sessions before a test session during which phasic dopamine was recorded. For this test session, rats were additionally presented with 14-15 trials in which each reward was presented in the absence of a cue (probe trials) and 14-15 trials in which a solenoid valve was activated without resulting in infusion to control for the possibility that rats were able to use the solenoid click as a predictor of sucrose delivery (dummy trials).

### Fast-scan cyclic voltammetry

On a single test day (behavioural component described above), phasic dopamine was recorded by lowering a carbon fiber microelectrode into NAc using a custom-built micromanipulator (UIC Research Resources Center). Procedures were similar to those described in detail elsewhere (Fortin et al., 2015). Briefly, a triangular voltage waveform (−0.4V → +1.3 V → −0.4 V relative to Ag/AgCl; 400 V/s; 10 Hz) was applied to the electrode using custom-built hardware (University of Washington Electromechanical Engineering). This results in the oxidation and reduction of electroactive species at the electrode surface. After subtracting the non-faradaic background signal, dopamine can be extracted from the data using principal component analysis (PCA). Training sets were derived in each rat by injecting a cocktail of cocaine (10 mg/kg) and raclopride (1 mg/kg) at the end of the recording session to evoke robust dopamine and pH changes. Electrodes were pre-calibrated in a custom flow cell (Sinkala et al., 2012) to derive a calibration factor. Across all electrodes, the average calibration factor was 45.55 nM/nA.

### Histology

Following completion of the experiment, rats were terminally anaesthetized with sodium pentobarbital (50 mg/kg). The position of each recording site was lesioned by lowering a polyimide-insulated stainless steel electrode to the same depth as the carbon fiber and passing current (4 x 4 s, 1 mA; Ugo-Basile Lesion Making Device) before rats were transcardially perfused with phosphate-buffered saline followed by neutral buffered formalin (10%). Fixed brains were sectioned on a cryostat (50 µm) and sections containing lesion marks were identified under light microscopy with assistance from a rat brain atlas (Paxinos and Watson, 1998).

### Data analysis and statistical methods

Behavioural data were acquired using Med-PC (Med Associates) and analyzed using custom MATLAB scripts. Voltammetry data were acquired and initially analyzed using TarHeel CV (Fortin et al., 2015) and then imported into Matlab (Mathworks) for further analysis. Statistical analysis was performed using SPSS (IBM) or Matlab. All analysis scripts and dopamine concentration data are available at https://github.com/mccutcheonlab/pellet-vs-intraoral. The raw data are available at doi:10.6084/m9.figshare.6387392. Probability of cue-evoked approach was defined as trials during which rats made a nose poke during the cue relative to total trials and was compared between cued pellet and cued infusion trials using Wilcoxon Signed Rank Test. Cue-evoked approach was defined as total time spent in food cup between cue onset and reward delivery and was compared between cued pellet trials and cued infusion trials using a paired t-test. Dopamine concentration traces were extracted from voltammetry data using principal component analysis (PCA). Trials in which the summed residual (Q) exceeded the threshold indicating satisfactory PCA (Qα) were automatically excluded (Keithley et al., 2009). Data were subsequently analyzed by comparing average dopamine concentration across different epochs. For uncued trials these epochs were: baseline (5 s before reward delivery) and reward (3 s following reward delivery). For cued trials these epochs were: baseline (5 s before cue onset), cue (1 s following cue onset), and reward (1 s following reward delivery). Epoch lengths were based on differences in pellet retrieval latency across trials and were designed to capture the large majority (>70%) of latencies. Two-way within-subjects ANOVA with Trial Type and Epoch as factors was used with appropriate *post hoc* tests (Bonferroni or Dunnett’s). Probability of cue-evoked approach and dopamine concentration during approach vs. no approach trials was analyzed with Wilcoxon Signed Rank Test. To probe regional differences, receiver operator characteristic (ROC) analysis was applied to dopamine concentration traces after they were binned into 500 ms. All trials from dopamine measurements in the core were compared to all trials from measurements in the shell for each trial type. p < 0.05 was considered as significance level.

## Author Contributions

JM designed experiments, carried out experiments, analyzed data and wrote the manuscript. MR designed experiments and wrote the manuscript.

## Funding Sources

This work was supported by the National Institutes of Health [grant numbers K01-DA033380 (JEM), R01-DA025634 (MFR)]; and the University of Illinois at Chicago Campus Research Board (JEM).

## Declaration of Conflicting Interests

The authors declare no conflicts of interest.

## Acknowledgements

We would like to thank Dr. Matthew Wanat for commenting on an earlier draft of the manuscript, Dr. Scott Ng-Evans for assistance with voltammetry components, and Eric Schmidt and the UIC Biological Resources Center for engineering and workshop help.

